# Unsupervised Topological Alignment for Single-Cell Multi-Omics Integration

**DOI:** 10.1101/2020.02.02.931394

**Authors:** Kai Cao, Xiangqi Bai, Yiguang Hong, Lin Wan

## Abstract

Single-cell multi-omics data provide a comprehensive molecular view of cells. However, single-cell multi-omics datasets consist of unpaired cells measured with distinct unmatched features across modalities, making data integration challenging. In this study, we present a novel algorithm, termed UnionCom, for the unsupervised topological alignment of single-cell multi-omics integration. UnionCom does not require any correspondence information, either among cells or among features. It first embeds the intrinsic low-dimensional structure of each single-cell dataset into a distance matrix of cells within the same dataset and then aligns the cells across single-cell multi-omics datasets by matching the distance matrices via a matrix optimization method. Finally, it projects the distinct unmatched features across single-cell datasets into a common embedding space for feature comparability of the aligned cells. To match the complex nonlinear geometrical distorted low-dimensional structures across datasets, UnionCom proposes and adjusts a global scaling parameter on distance matrices for aligning similar topological structures. It does not require one-to-one correspondence among cells across datasets, and it can accommodate samples with dataset-specific cell types. UnionCom outperforms state-of-the-art methods on both simulated and real single-cell multi-omics datasets. UnionCom is robust to parameter choices, as well as subsampling of features. UnionCom software is available at https://github.com/caokai1073/UnionCom.

## 1 Introduction

The advent of single-cell sequencing provides high-resolution omics profiles at cellular level (e.g. single-cell DNA sequencing (scDNA-seq) of genomics, single-cell RNA sequencing (scRNA-seq) of transcriptomes, single-cell ATAC-sequencing (scATAC-seq) of chromatin accessibility), offering the opportunity to unveil molecular mechanisms related to fundamental biological questions, such as cell fate decisions [1]. Although extensive studies have been conducted on the computational analysis of single-cell genomic, transcriptomic, and epigenetic data, most methods are limited to handling single-cell data in one modality, yielding a series of separated and patched views of the intrinsic biological processes. Recently, single-cell multi-omics data across modalities profiled from cells sampled from the same sample or tissue are emerging. Integration of single-cell multi-omics data will build connections across modalities, providing a much more comprehensive molecular multi-view of intrinsic biological processes [2, 3].

The integration of single-cell omics datasets is drawing heavy attention on advances in machinelearning and data science [2, 3]. Many single-cell data integration methods have been developed for the batch-effect correction of single-cell datasets in one modality [2]. However, compared with the single-cell batch-effect correction problem, the integration of single-cell multi-omics datasets across modalities poses fresh challenges in two ways. First, single-cell multi-omics datasets consist of unpaired cells measured with distinct unmatched features across modalities [2]. As most single-cell sequencing assays are still destructive for cells, single-cell datasets of either the same modality or different modalities generally have unpaired cells. Moreover, single-cell multi-omics datasets do not share common features across modalities, since each modality is aimed at acquisition of cellular molecular identity from a particular aspect. The distinct features across modalities reside in different mathematical spaces. Generally, therefore, single-cell multi-omics data do not have any correspondences, either among samples (cells) or among features. In practice, empirical pre-matching of distinct features across modalities into a common space based on *prior* knowledge is generally taken prior to applying state-of-the-art single-cell data integration methods. Second, single-cell multi-omics data are generated by different assays, with distinct underlying data generative models and mechanisms. Therefore, complex nonlinear geometrical distortions on datasets across modalities can be introduced to the biological intrinsic low-dimensional structures. Such complex distortions make the integration of single-cell multi-omics data much more difficult than batch-effect correction for single-cell datasets in one modality. Therefore, linear operations, such as translation and scaling, which generally work well for batch-effect correction of single-cell datasets in one modality, are now inadequate for the alignment of single-cell multi-omics datasets with complex nonlinear geometrical distortions across modalities.

Computational methods have been developed for single-cell data integration. However, many existing methods were designed for batch-effect correction [2]. The challenges of single-cell multi-omics integration arise as a result of distinct unmatched features, as well as complex nonlinear geometrical distortions across modalities, and these problems remain unresolved. The pioneering single-cell data integration method, MNNCorrect [4], aimed to remove batch effects across datasets by aligning mutual nearest neighbors (MNNs), as calculated by the Euclidean distances between cells across datasets in a common feature space. However, MNNCorrect is restricted to cases in which batch-effect is almost orthogonal to the biological subspace and cases in which batch-effect variation is much smaller than biological-effect variation between different cell types [4]. The established single-cell data analysis pipeline, Seurat [5], projected (distinct) feature spaces across datasets using canonical correlation analysis (CCA) into a common subspace which maximizes the inter-dataset correlation structure, and then adopted a strategy similar to that of MNNCorrect to align the cells between datasets by finding anchor cells (i.e. corresponding points) across datasets based on the MNNs calculated from the projected common subspace. However, since CCA is a linear dimensionality reduction method, it cannot capture the nonlinear interrelationships among singlecell multi-omics data across modalities. Scanorama [6] was built upon the MNN-based strategy for the computationally efficient integration of many large-scale scRNA-seq datasets. ScAlign [7] developed a deep autoencoder-based approach for the nonlinear dimensionality reduction of shared common feature space across one-modality scRNA-seq datasets and then aligned cells across datasets.

Methods developed for single-cell multi-omics integration across modalities are emerging. Liger [8], which worked on datasets with pre-matched common feature space across modalities, employed non-negative matrix factorization to find the shared low-dimension factors of the common features to match single-cell omics datasets. MATCHER [9] removed the requirement for correspondence information of features across modalities and integrated single-cell multi-omics datasets based on manifold alignment. However, MATCHER is limited to the alignment of 1-D trajectories [9], but it is incapable of aligning complex trajectories, such as branched tree structures. MAGAN [10] integrated single-cell multi-omics datasets by aligning the biological manifolds via a generative adversarial networks (GANs) approach. However, it lacked power in unsupervised task when the correspondence information among samples across datasets is unavailable [11]. Harmonic [12] aligned the diffusion geometries across datasets based on diffusion map method, but it required partial feature correspondence information. MMD-MA [11], an unsupervised manifold alignment algorithm for single-cell multi-omics datasets, embeds the latent biological low-dimensional structures in Reproducing Kernel Hilbert spaces (RKHSs), and found a shared common subspace of the RKHSs for manifold alignment by minimizing the maximum mean discrepancy (MMD) across modalities.

In the present study, we present a novel algorithm, termed UnionCom, for the unsupervised topological alignment of single-cell multi-omics integration. UnionCom is an unsupervised method which does not require any correspondence information, either among cells or among features, across single-cell multi-omics datasets. It extends the generalized unsupervised manifold alignment (GUMA) algorithm [13], which was originally applied to the 3-D structure alignment of protein sequences, to tackle the difficulty in the topological alignment of complex nonlinear distorted intrinsic low-dimensional structures embedded in the single-cell data. Specifically, UnionCom first embeds the intrinsic low-dimensional structure of each single-cell dataset into a distance matrix of cells within the same dataset and then aligns the cells across single-cell multi-omics datasets by matching the distance matrices via a matrix optimization method. Finally, it projects the distinct unmatched features across single-cell datasets into a common embedding space for feature comparability of the aligned cells (Fig. 1). UnionCom works on the general assumption that cells of single-cell multi-omics datasets are from similar cell populations sampled from similar biological processes or tissues with similar intrinsic low-dimensional structures embedded in the data. It does not require one-to-one correspondence among cells across datasets, and it can take care of samples with dataset-specific cell types. We demonstrate the power of UnionCom in unsupervised topological alignment for single-cell multi-omics integration, especially for complex intrinsic structure embeddings, using both simulated and real single-cell multi-omics datasets. Compared with state-of-the-art methods, UnionCom outperforms them with high accuracy. In addition, UnionCom is robust in terms of parameter choices, as well as subsampling of features.

**Figure 1:**
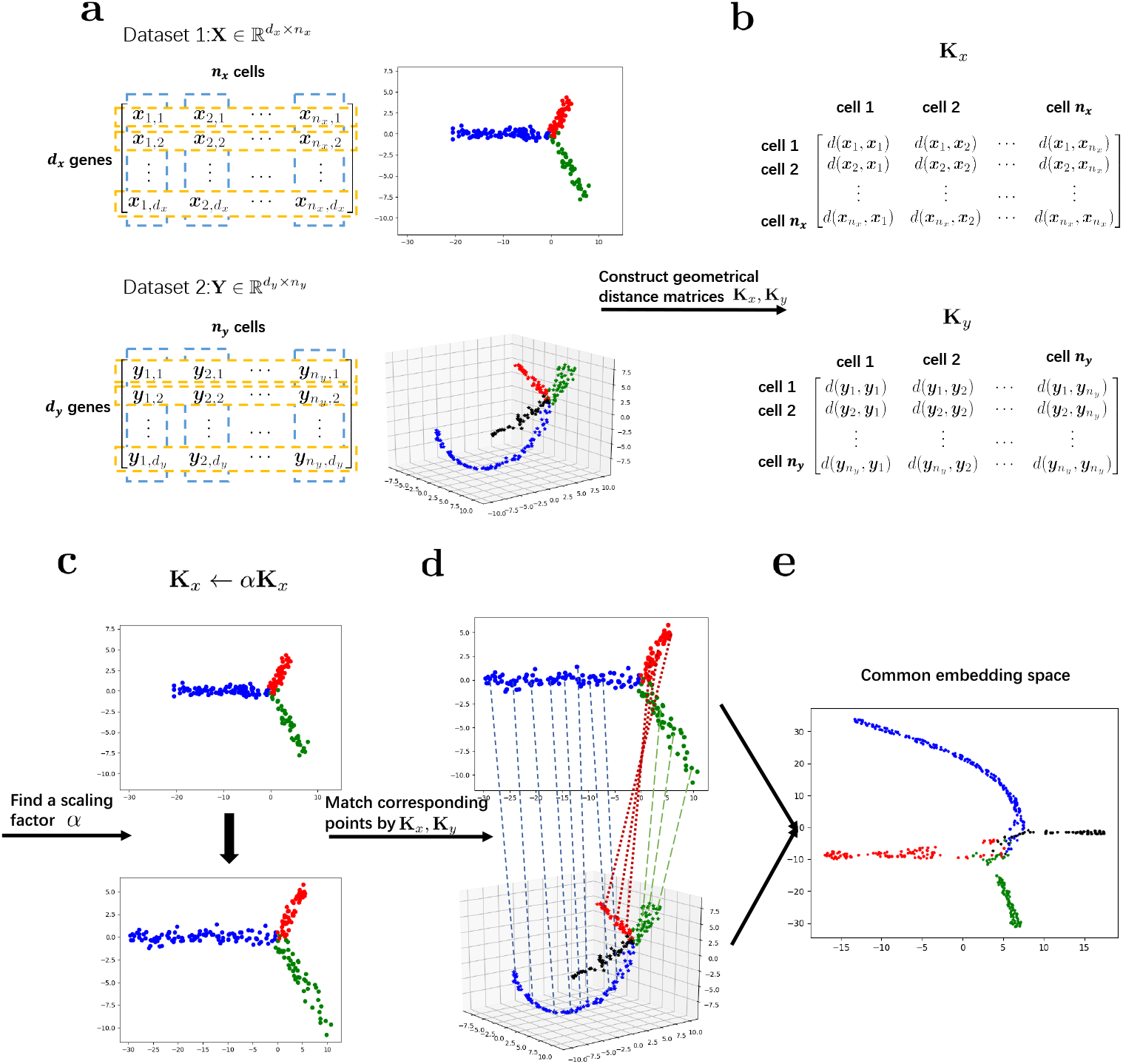
Schematic overview of UnionCom. (a) Given the input of single-cell multi-omics datasets (e.g., Dataset 1 and Dataset 2), which have similar embedded topological structures, UnionCom (b) embeds the intrinsic low-dimensional structure of each single-cell dataset into a geometrical distance matrix of cells within the same dataset; (c) rescales the global distortions on the topological structures across datasets by a global scaling parameter *α*; (d) aligns the cells across single-cell datasets by matching the geometrical distance matrices based on a matrix optimization method; and (e) finally projects the distinct unmatched features across modalities into a common embedding space for feature comparability of the aligned cells. It does not require one-to-one correspondence among cells across datasets, and it can accommodate samples with dataset-specific cell types (see the branch with black points in Dataset 2 for example).

## 2 Method

### 2.1 UnionCom Algorithm

UnionCom is an unsupervised topological alignment algorithm for single-cell multi-omics integration (Fig. 1). We describe details of UnionCom as follows.

Here, we formulate our method for the case of two datasets. However, it can be easily generated for cases of any number of single-cell multi-omics datasets. Suppose we have two single-cell multi-omics datasets, 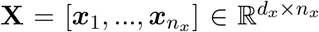 and 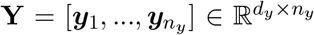, across two modalities where *d_x_*(*d_y_*) and *n_x_*(*n_y_*) are the number of features (e.g., gene expression, DNA methylation) and cells for the **X** (**Y**), respectively. Without loss of generality, we assume that *n_x_ ≤ n_y_*. We assume that the intrinsic low-dimensional manifolds of **X** and **Y** are 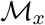 and 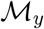, respectively, which share similar topological structure. Given the input of **X** and **Y**, UnionCom aligns the cells between the datasets and then projects the distinct unmatched features into a common embedding space as the coordinates of the aligned cells between datasets. UnionCom consists of the following three major steps (A1-A3).

#### A1. Embedding the intrinsic low-dimensional structure of each single-cell dataset into the geometrical distance matrix

UnionCom embeds the intrinsic low-dimensional structure of each single-cell dataset into a metric space by using the geometrical distance matrix of the cells within the same dataset which is defined as [**K**]_*ij=d*_(***x**_i_*, ***x**_j_*), where *d* is a geodesic distance between cells on the intrinsic manifold. Accordingly, **K**_*x*_ and **K**_*y*_ represent geometrical distance matrices for 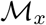 and 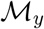, respectively. To calculate the geodesic distance, UnionCom first constructs a *k* nearest neighbor graph (*k*-nn graph) of cells for each dataset and then calculates the shortest distance between each pair of nodes (cells) on the graph using the Floyd-Warshall algorithm since the shortest distance path will approximate to geodesic distance [14]. We set *k* to be the minimum number that makes *k*-nn graph connected. We demonstrate that UnionCom is robust to the choices of *k* in a wide range (Fig. 5(b)).

#### A2. Aligning cells across single-cell multi-omics datasets by matching the geometrical distance matrices

UnionCom aligns cells across single-cell datasets by matching the geometrical distance matrices. By extending the unsupervised manifold alignment algorithm GUMA [13], UnionCom proposes and adjusts a global scaling parameter *α* in Eq. (1) for rescaling the global geometrical distortions for the topological structure alignment across datasets. In addition, UnionCom relaxes the restriction of one-to-one cell correspondence required by GUMA. It is capable of handling samples with dataset-specific cell types.

Specifically, UnionCom formalizes the unsupervised topological alignment problem into a matrix optimization problem as follows:

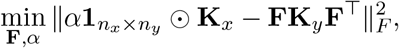

where **F** = [*F_ij_*]_*n_x_ × n_y_*_ is a point matching matrix satisfying certain constraints needed for matching; the 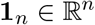 and 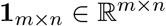 represent vector and matrix of ones; the superscript ⊤ denotes the transpose of a vector or matrix, and 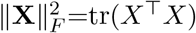 is the Frobenius norm (tr(·) is the trace norm). The *α*, which is newly proposed by UnionCom, is a global scaling parameter for the rescaling of the global geometrical distortions between datasets; the *α**1**_n_x_×n_y__* stands for a matrix with elements all of *α*; and the ⊙ denotes Hadamard product which is the element-wise product taken on two matrices of the same dimensions.

In the GUMA algorithm, **F** ∈ {0,1}^*n_x_×n_y_*^ is a hard matching matrix, i.e., a 0-1 integer matrix, to mark the correspondences of cells between **X** and **Y**, where [**F**]_*ij*_ = 1 denotes that ***x**_i_* and ***y**_j_* are the counterpart of each other. Therefore, GUMA implicitly assumes that individual cells are matched in a one-to-one fashion between the two datasets, a phenomenon that is often inconsistent with real data. GUMA finds the **F** over the constraint set Π of all possible 0-1 integer matrices

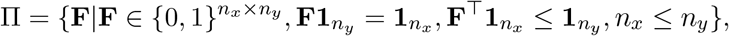

using KuhnC-Munkres algorithm. However, the set Π is neither close nor convex, leading to an NP-hard optimization problem.

UnionCom seeks a soft matching of cells between datasets. It relaxes the constraints of **F** to the following compact and convex set Π′ as

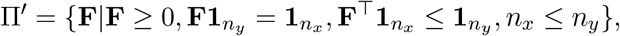

where **F** ≥ 0 denotes that each element in the matrix is non-negative and that the summation of each row of **F** equals 1. The **F** ∈ Π′ provides a probabilistic interpretation of sample matching between two datasets: [**F**]_*ij*_ indicates the likelihood of ***x**_i_* matching with ***y**_ij_*. Therefore, UnionCom aligns the topological structures between two datasets by solving the matrix optimization problem over the constraint set Π′ as follows:

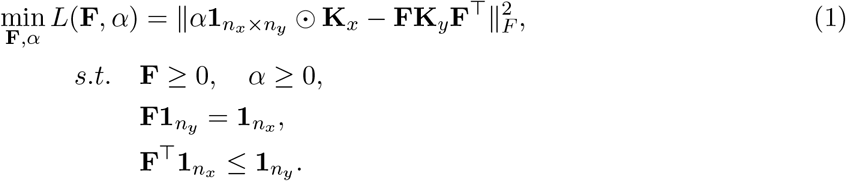

We develop a prime-dual method to solve the optimization problem of Eq. (1) by minimizing an augmented Lagrangian function as

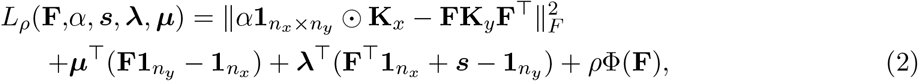

where ***μ*** and **λ** are the Lagrangian multipliers; the *n_y_* dimensional non-negative vector ***s*** ≥ 0 is a slack variable that transforms the inequality constraint to an equality constraint of **F**^⊤^**1**_*n_x_*_+***s*** = **1**_*n_y_*_; 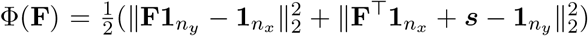 is the penalty term added to make the solution stable and converging fast (Fig. 5(d)); and *ρ* is a user-defined hyperparameter. UnionCom is robust for the choice of *ρ* (Fig. 5(d)), and we set its default as *ρ* = 10.

The proposed prime-dual algorithm minimizes (2) over **F** ∈ Π′ and *α* by the following iterative steps

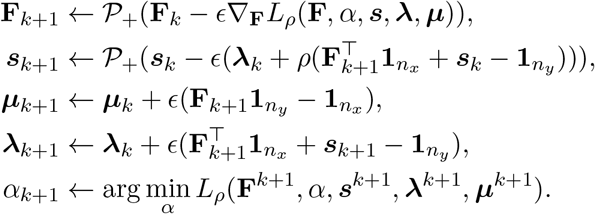

where 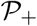 is a projection operator that maps the function into the non-negative quadrant. The *ϵ* is the learning rate which is often set as a small value (in generally less than 1*e* − 3). By alternating iterations on the scaling factor *α* and matching matrix **F**, UnionCom reaches an optimal solution over the relaxed convex constraint set Π′. For numerical stability, we normalize **K**_*x*_ ← **K**_*x*_/ max(*n_x_, n_y_*) and **K**_*y*_ ← **K**_*y*_/ max(*n_x_, n_y_*) before applying prime-dual algorithm. See the pseudocode and the derived gradients on the website of UnionCom software.

It is worth noting that UnionCom can be easily extended to supervised or semi-supervised alignment by fixing the corresponding elements of **F** to be 1 when correspondence information of cells between datasets is available.

#### A3. Projecting distinct unmatched features across single-cell multi-omics datasets into a common embedding space

UnionCom projects the distinct unmatched features across single-cell multi-omics datasets into a common embedding space as the coordinates for the aligned cells across datasets. To preserve both the intrinsic low-dimensional structures and the aligned cells together simultaneously, UnionCom builds upon a t-Distribution Stochastic Neighbor Embedding (t-SNE) method as in [15].

Specifically, UnionCom aims to embed the datasets 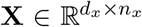 and 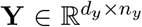 into a common embedded space of *p*-dimension (*p* ≤ min{*d_x_,d_y_*}), with the dimensionality reduced datasets denoted as 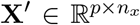 and 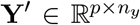, respectively. UnionCom finds the optimal 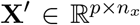 and 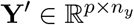 by minimizing a loss function 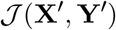 as follows:

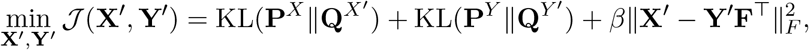

where **F** is the optimal matching matrix obtained in step A2; 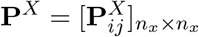 and 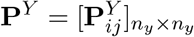 are the cell-to-cell transition probability matrix defined in the original spaces of **X** and **Y**, respectively, as in t-SNE; and 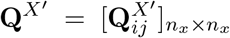 and 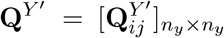, which are constrained to t-distribution, are the cell-to-cell transition probability matrix defined in the dimensionality reduced common space of **X**′ and **Y**′, respectively, as in t-SNE; the KL-divergence is defined as 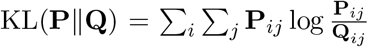. The loss function 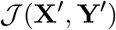 has three items: two KL-divergence terms between **P**^*X*^(**P**^*Y*^) and **Q**^*X′*^ (**Q**^Y*′*^) for the two datasets, respectively, and a coupling term for measuring the distance between the matched datasets in the common embedding space. The *β* is a trade-off parameter to balance the two KL terms and the coupling term. UnionCom uses the *ℓ*_2_ norm to measure the distance of matched datasets, but other distances in the embedding space can also be used. We solve this optimization problem by the gradient descent method. See the website of UnionCom software for more details. We set *p* = 32 as defaults when the size of the features is above 100.

It is worth noting that, UnionCom is not restricted to the t-SNE-based approach for step A3. Dimensionality reduction methods, such as elastic embedding [16] and UMAP [17, 18], both of which preserve local and global structures of data, can also be adopted by UnionCom for step A3.

### 2.2 Data

#### 2.2.1 Simulated Data

We simulate two sets of single-cell multi-omics datasets as follows: 1) Simulation 1 contains two datasets, which share a similar complex tree with two branching points embedded in distinct 2-D spaces (Fig. 2(a), left panels); 2) Simulation 2 contains two datasets: Dataset 1 has a bifurcated tree embedded in 2-D space (Fig. 2(b), upper left panel), while Dataset 2 has a trifurcated tree embedded in 3-D space with one branch as the dataset-specific cell type (Fig. 2(b), lower left panel; the green branch is the dataset-specific cell type). The correspondence information of cells between the two datasets within the same simulation is known from simulation. We project the 2 simulated datasets into high-dimensional feature spaces (with dimensionality of 1000 for Dataset 1 and 500 for Dataset 2, respectively, for each simulation) by adding the feature vectors with elements being sampled randomly from Gaussian distribution.

**Figure 2:**
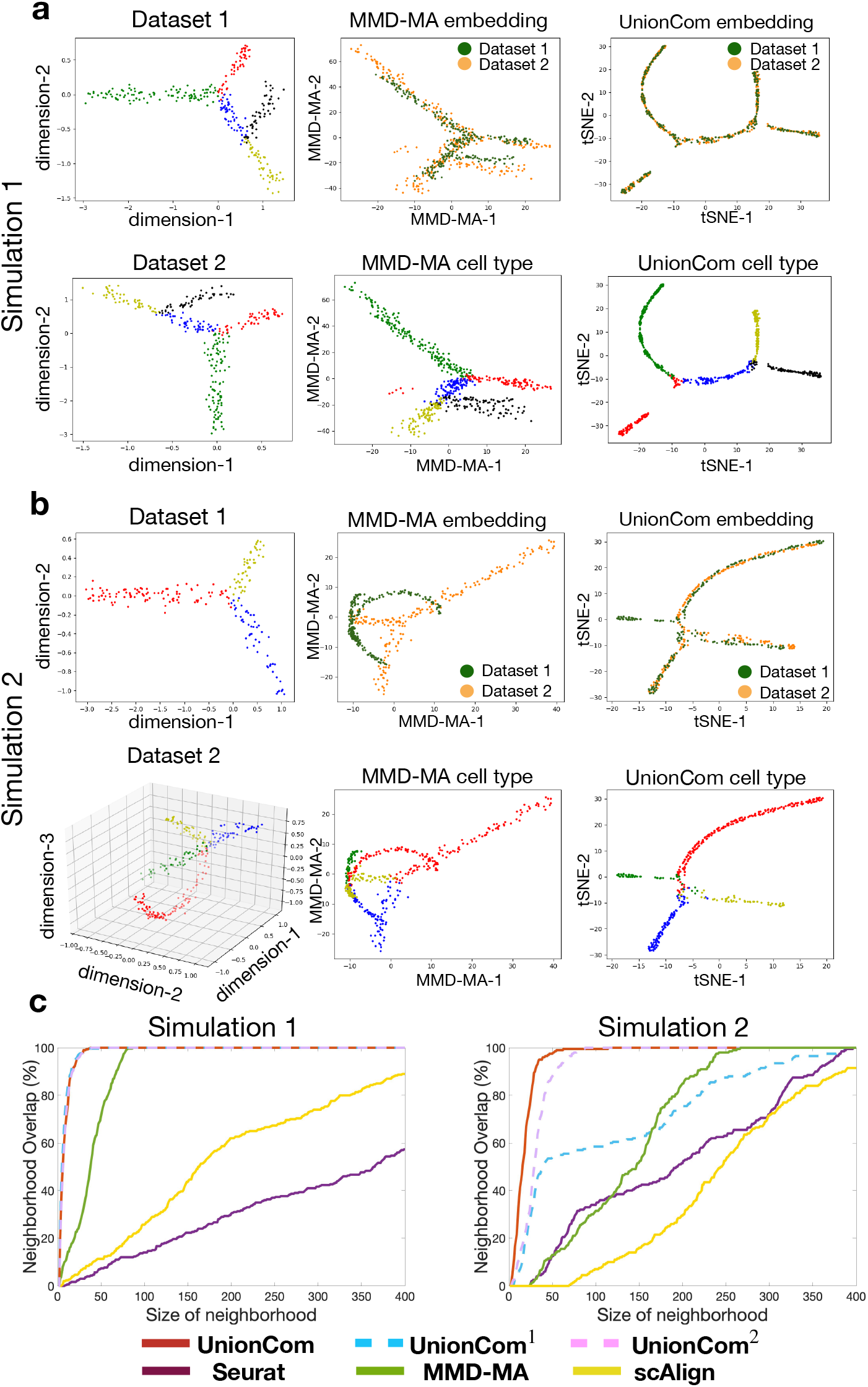
Alignment of simulated datasets. (a) Simulation 1 and (b) Simulation 2: Visualizations of Dataset 1 (upper left panel) and Dataset 2 (lower left panel), separately using t-SNE before alignment; branches with points in the same colors are matched between datasets; visualization of the common embedding space of the two aligned datasets by MMD-MA (upper middle panel: points are colored according to their corresponding datasets; lower middle panel: points are colored according to their corresponding branches) and UnionCom (upper right panel: points are colored according to their corresponding datasets; lower right panel: points are colored according to their corresponding branches). The green branch of Dataset 2 of Simulation 2 (lower left panel of (b)) is a dataset-specific cell type. (c) Averaged percentage of Neighborhood Overlap at different size of neighborhood (left panel: Simulation 1; right panel: Simulation 2).

#### 2.2.2 Real single-cell multi-omics datasets

We use two real sets of single-cell multi-omics of sc-GEM data [19] and sc-NMT data [20]. The first real set generated by sc-GEM sequencing, hereinafter denoted as sc-GEM data, contains two single-cell omics datasets of gene expression and DNA methylation on samples from human cells undergoing reprogramming to iPSCs. The data were generated by [19] and used in MATCHER [9]. The second real data set generated by sc-NMT sequencing, hereinafter denoted as sc-NMT data, contains three single-cell omics datasets of gene expression, DNA methylation, and chromatin accessibility on samples from mouse gastrulation collected at 3 time stages (embryonic day 5.5 (E5.5), E6.5, and E7.5). The data were generated by [20].

The sc-GEM sequencing measured the gene expression and DNA methylation of the same cell simultaneously; and the sc-NMT sequencing measured the gene expression, DNA methylation, and chromatin accessibility of the same cell-cell simultaneously. Thus, for each of the real data sets, the cell correspondence information across single-cell multi-omics datasets is available.

### 2.3 Method Evaluations

We evaluate the single-cell multi-omics integration methods using two indexes, 1) Neighborhood Overlap and 2) Label Transfer Accuracy, to measure the alignment accuracies. Both indexes work on the basis of the common embedded space (coordinate) of the integrated datasets.

When the cell-cell correspondence information between multi-omics datasets is available, the Neighborhood Overlap, which was proposed by Harmonic [12], is used to measure the ability to recover the one-to-one correspondence of cells between two datasets: for a given size of neighborhood of each cell in the common embedded space, the percentage of neighborhood overlap of a dataset is defined as the percentage of cells that can find their correspondence cells from the other dataset in their neighborhood, respectively. We use the averaged percentage of neighborhood overlap of the two datasets. The averaged percentage of neighborhood overlap ranges from 0 to 100%, and a higher percentage is indicative of a better recovery of cell-to-cell relationship between two datasets.

When the cell label information (e.g., cell types, branches of cell trajectories) is available, Label Transfer Accuracy, which has been widely used in the transfer learning community and was adopted by scAlign [7], is used to measure the ability of transferring labels of cells from one dataset to another in the common embedded space. Assuming that Dataset 2 has more cells than Dataset 1, we use Dataset 2 as the training set and Dataset 1 as the testing set. We construct a *k^acc^*-nn classifier trained by cells with their labels using Dataset 2, and Label Transfer Accuracy is the prediction accuracy of the cell labels on the testing set, i.e., Dataset 1. The value of Label Transfer Accuracy, which is the percentage of cells with correctly predicted labels among all predicted cells, ranges from 0 to 100%, and a higher percentage is indicative of a better performance in transferred labels based on the two aligned datasets. We set *k^acc^* = 5 as default, but the Label Transfer Accuracy is stable across different choices of *k^acc^* (Fig. 3(f)).

**Figure 3:**
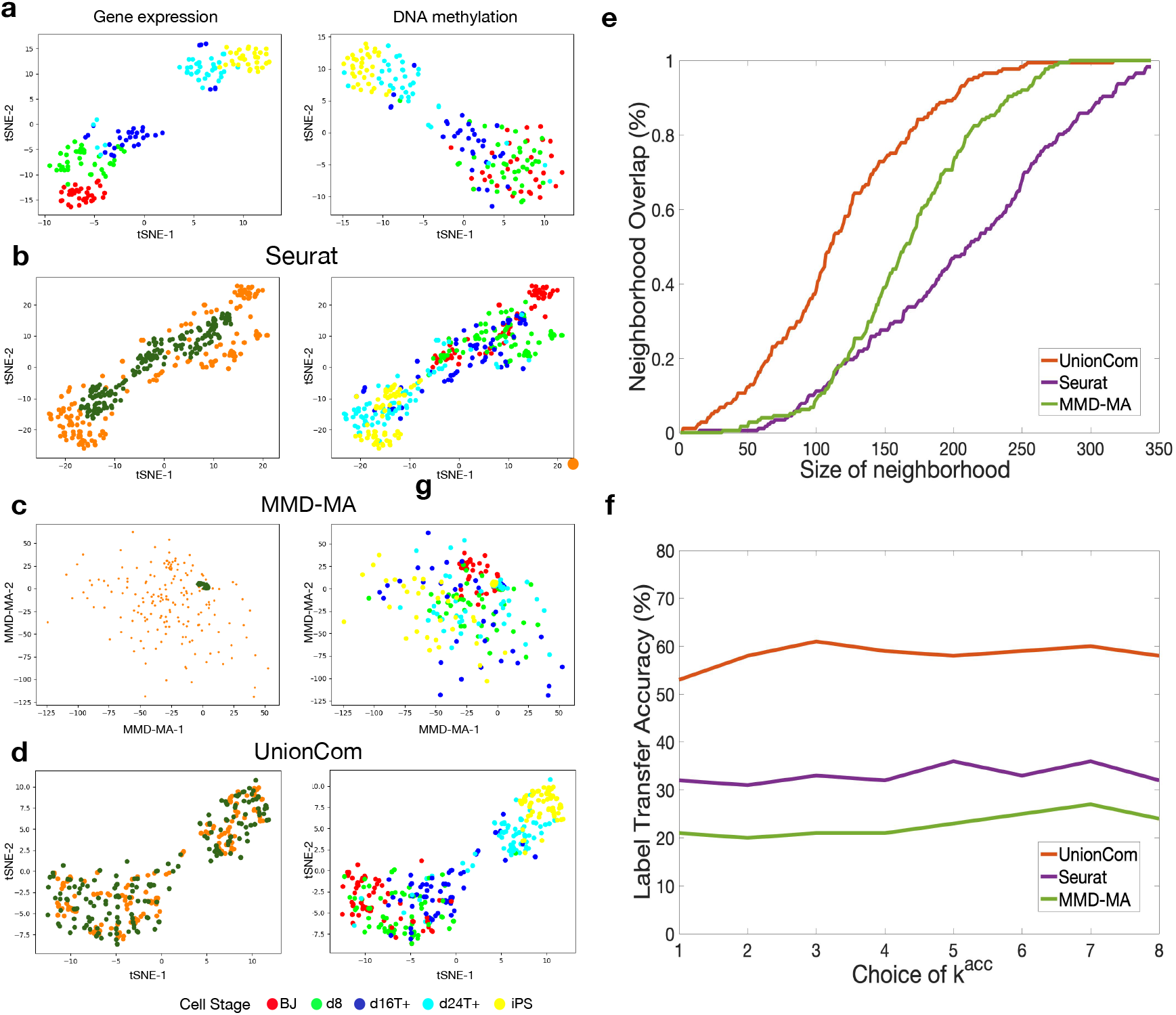
Alignment of sc-GEM omics datasets of gene expression and DNA methylation. (a) Visualizations of the gene expression and DNA methylation datasets separately using t-SNE before alignment. (b,c,d) Visualizations of the common embedding space of the two aligned datasets by Seurat (b), MMD-MA (c) and UnionCom (d), respectively (left panel: points (cells) are colored according to their corresponding datasets; right panel: points (cells) are colored according to their corresponding cell types). (e) Averaged percentage of Neighborhood Overlap at different size of neighborhood. (f) Label Transfer Accuracy at different *k^acc^* of the *k^acc^*-nn classifier.

## 3 Results

### 3.1 UnionCom outperforms the state-of-the-art methods on integrating simulated datasets with highest accuracies

We simulate two sets of single-cell multi-omics datasets. Each simulated set has similar complex topological structures, but with different geometrical distortions embedded in distinct high-dimensional spaces (Section 2.2).

We apply UnionCom for the alignment of the simulated data sets. We project the common space of the two aligned datasets by UnionCom to a 2-D space for visualization using t-SNE. For each of the simulations, UnionCom integrates the two datasets with well-aligned geometrical structures (see upper right panels of Fig. 2(a,b)) and well-matched branches (see lower right panels of Fig. 2(a,b)). Although Dataset 2 of Simulation 2 has a dataset-specific branch (see the branch with green-colored points shown in the lower left panel of Fig. 2(b)), UnionCom still shows its accuracy in aligning the two datasets accordingly, leaving the dataset-specific branch unaligned to any other branches (see the lower right panel of Fig. 2(b)). For our comparisons, we also apply the MMD-MA algorithm, which adopts a MMD term to reduce distribution discrepancy in feature spaces and to align the simulated datasets using its defaulting parameters. Although MMD-MA can align the two datasets relatively well in Simulation 1 (Fig. 2(a), upper middle panel), it fails to merge samples from the two datasets of Simulation 2 into common regions (Fig. 2(b), upper middle panel).

We further evaluate the accuracy of the aligned datasets using indexes of both Neighborhood Overlap and Label Transfer Accuracy (Section 2.3) and compare the performances of UnionCom with that of the state-of-the-art methods Seurat v3, scAlign and MMD-MA. In addition, to validate the effectiveness of using geodesic distance and global scaling factor *α*, we also test UnionCom using Euclidean distance instead of geodesic distance (denoted as UnionCom^1^), as well as UnionCom with a fixed 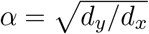 (denoted as UnionCom^2^) for our comparisons. We do not include MAGAN and Harmonic in our comparisons because both need correspondence information, either among samples or among features between datasets.

Among the 6 methods tested, UnionCom always has the highest accuracy in both Neighborhood Overlap and Label Transfer Accuracy on the two simulated datasets (Fig. 2(c) and Table 1). UnionCom^1^, which uses Euclidean distance, achieves almost the same accuracy as that of UnionCom in Simulation 1, but dramatically drops in accuracy on Simulation 2 in which the embedded manifolds are nonlinearly distorted. UnionCom^2^ does not incorporate the global scaling parameter *α*, but it still ranks third and second highest in accuracy for Simulations 1 and 2, respectively. MMD-MA has moderate accuracy performance on both Simulations 1 and 2 (Fig. 2(c) and Table 1). It is also not surprising that scAlign and Seurat v3 do not achieve high accuracy since both were mainly developed for integrating scRNA-seq datasets of one modality.

**Table 1:**
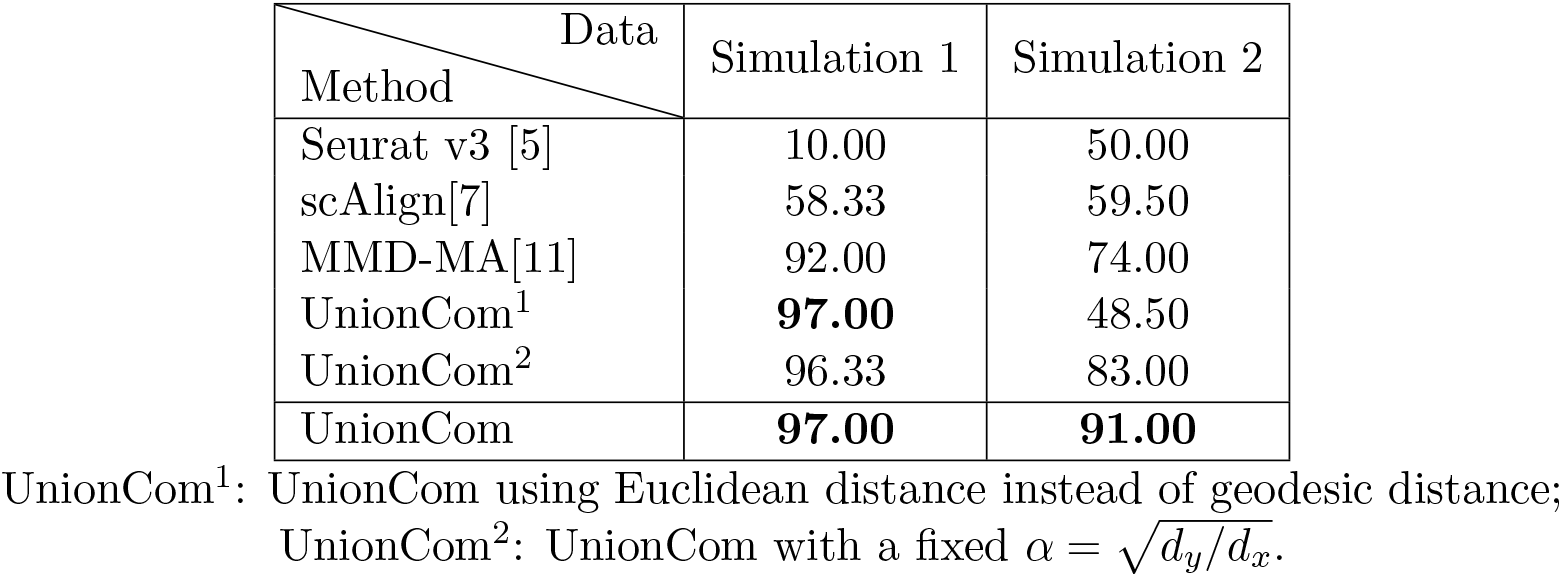
The Label Transfer Accuracy (%) by 6 methods on the 2 simulation studies.

### 3.2 UnionCom accurately integrates sc-GEM omics datasets across 2 modalities

We apply UnionCom to integrate sc-GEM omics datasets of gene expression and DNA methylation (see Section 2.2 for details). We include Seurat v3, which has a CCA step, to find a common feature space across datasets, and MMD-MA which is unsupervised, for our comparisons. We obtain the processed sc-GEM data from MATCHER [9], which contain 177 cells with 34 features in the gene expression dataset and 177 cells with 27 features in the DNA methylation dataset. The cell-type annotation is from [19].

We visualize the gene expression dataset and the DNA methylation dataset separately using t-SNE with cells being colored by their annotated cell types before alignment (Fig. 3(a)). Both datasets demonstrate similar linear structures with the same orders on cell types: the BJ cells (red points) locate at one end, and the iPS cells (yellow points) locate at the other end of the linear trajectories. This is consistent with the underlying processes of cells undergoing reprogramming to iPSCs.

Since sc-GEM data have relatively small feature sizes, we embed the two datasets in a common space using UnionCom with dimensionality of *p* = 2 and visualize them on the 2-D UnionCom space (Fig. 3(d)). We find that UnionCom aligns the cells between the two datasets quite well by locating samples between datasets on a common region with similar distributions (Fig. 3(d), left panel). In contrast, Seurat v3 locates cells from the gene expression dataset in an interior region surrounded by cells from the DNA methylation dataset outside (Fig. 3(b), left panel); and MMD-MA does not put the cells between two datasets on comparable scales, because the cells from the gene expression dataset are all collapsed together (Fig. 3(c), left panel). When looking at the cell-type labels, we find that UnionCom (Fig. 3(d), right panel) has the best separation of cell types as well as preservation of the global structures of cell lineage on the merged datasets compared to Seurat v3 (Fig. 3(b), right panel) and MMD-MA (Fig. 3(c), right panel).

In addition, UnionCom achieves highest accuracy in both Neighborhood overlap (Fig. 3(e)) and Label Transfer Accuracy (Fig. 3(f)) when compared with Seurat v3 and MMD-MA.

### 3.3 UnionCom accurately integrates sc-NMT omics datasets across 3 modalities

We obtain the sc-NMT data from [20] (see Section 2.2 for details). We filter out cells with all features being denoted as missing values (“NA”) for each of the 3 datasets, resulting in 1940 cells with 5000 features in the gene expression dataset, 709 cells with 2500 features in the DNA methylation dataset, and 612 cells with 2500 features in the chromatin accessibility dataset, respectively. We find that UMAP [17, 18] can obtain a best preservation of the global structure of cell lineage for the sc-NMT data than PCA and t-SNE (Fig. 4(a)). Therefore, we apply UMAP to conduct the dimensionality reduction of each of the 3 datasets to a dimensionality of 300, respectively, prior to the alignment using UnionCom, Seurat v3 and MMD-MA.

**Figure 4:**
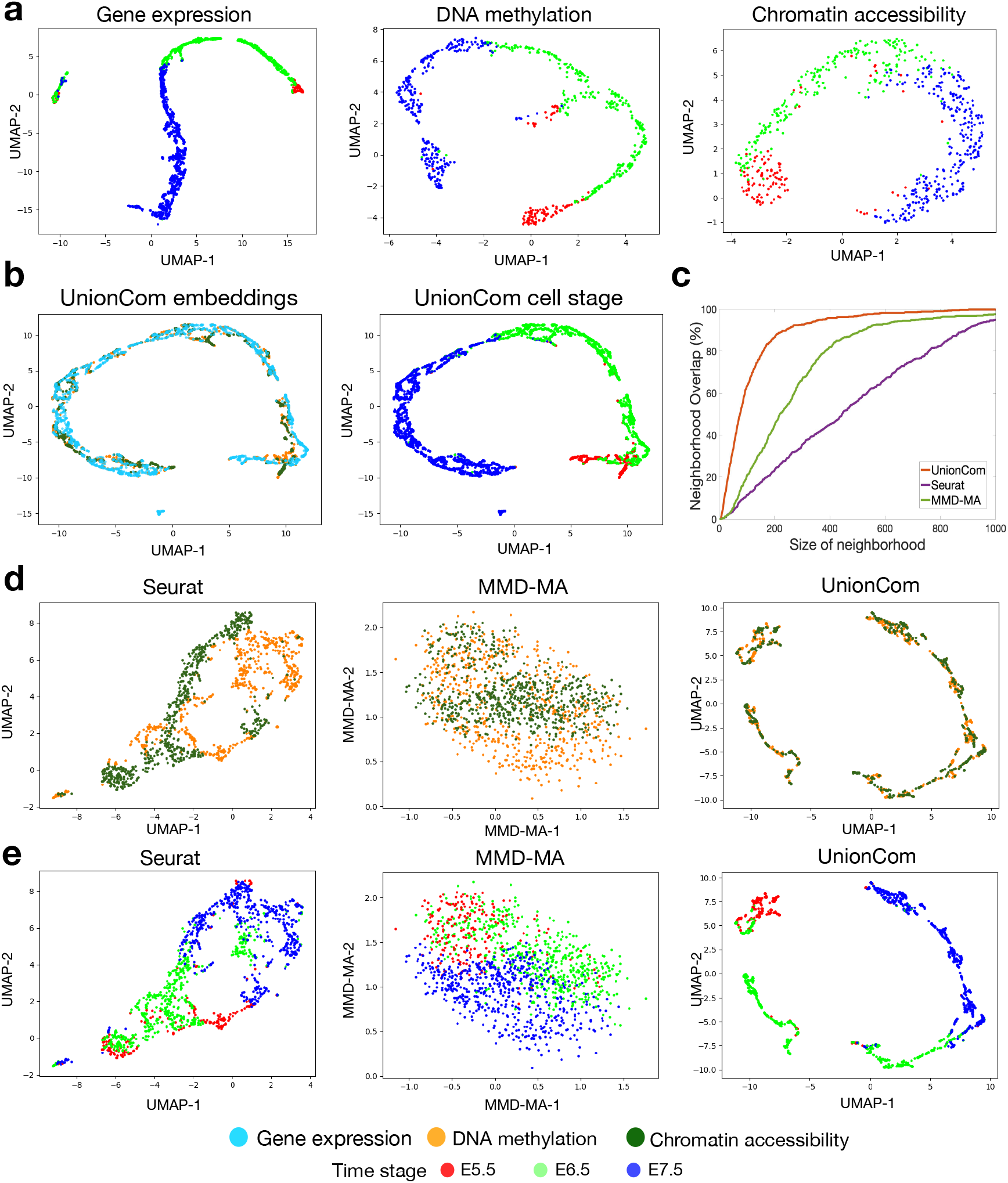
Alignment of sc-NMT omics datasets of gene expression, DNA methylation, and chromatin accessibility. (a) Visualizations of the gene expression, DNA methylation, and chromatin accessibility datasets separately using UMAP before alignment. (b) Visualizations of the common embedding space of the three datasets aligned by UnionCom (left panel: points (cells) are colored according to their corresponding datasets; right panel: points (cells) are colored according to their corresponding time stages). (c) Averaged percentage of Neighborhood Overlap on the alignment of the two datasets of DNA methylation and chromatin accessibility. (d,e) Visualizations of the common embedding space of the two aligned datasets of DNA methylation and chromatin accessibility by Seurat (left panel), MMD-MA (middle panel) and UnionCom (right panel), respectively; in (d): points (cells) are colored according to their corresponding datasets; in (e): points (cells) are colored according to their corresponding time stages).

We visualize the three datasets of gene expression, DNA methylation and chromatin accessibility separately using UMAP with cells being colored by their annotated time stage (e.g., E5.5, E6.5 and E7.5) (Fig. 4(a)). All 3 datasets demonstrate similar dominant linear structures of cell lineage, consistent with the underlying processes of cell development during mouse gastrulation from E5.5 to E7.5.

We apply UnionCom to integrate sc-NMT omics datasets of gene expression, DNA methylation, and chromatin accessibility simultaneously and embed them into a common space (Fig. 4(b)). For multiple datasets, UnionCom selects the dataset with largest sample size as the reference dataset and aligns the other datasets with respect to the reference. When visualizing the common embedded space of the three datasets by UnionCom using UMAP in 2-D space, we find that UnionCom integrates the 3 datasets across modalities quite well by aligning cells across datasets along a dominant linear trajectory on similar regions (Fig. 4(b), left panel), and also by preserving the global structure of time stage orders (Fig. 4(b), right panel).

In addition, we include Seurat v3 and MMD-MA as our comparisons and demonstrate their performances on the alignment of the two datasets of DNA methylation and chromatin accessibility. UnionCom still achieves highest accuracy in both Neighborhood Overlap (Fig. 4(c)) and Label Transfer Accuracy when using time stages as cell labels (Label Transfer Accuracy: UnionCom: 85.95%, MMD-MA: 45.65% and Seurat 53.27%). We find that UnionCom aligns the cells between the two datasets quite well in 2-D space by aligning the cells between the datasets along a linear trajectory and by merging the two datasets on a common region with similar distributions (Fig. 4(d), right panel). In contrast, both Seurat v3 (Fig. 4(d), left panel) and MMD-MA (Fig. 4(d), middle panel) do not merge the cells between two datasets into comparable spaces. When looking at the cell labels of time stages, we find that UnionCom preserves the global structures of time stage orders better than Seurat v3 and MMD-MA (Fig. 4(e)).

### 3.4 Robustness analysis of UnionCom

We confirm the robustness of UnionCom in both subsampling features of data and parameter choices on the two scNMT datasets for DNA methylation and chromatin accessibility. When randomly sampling a subset of features without replacement from both DNA accessibility and chromatin methylation datasets separately prior to data alignment, the label transfer accuracies by UnionCom are stable, showing much smaller fluctuation than those by Seurat and MMD-MA (Fig. 5(a)). When choosing *k* of the *k*-nn graph in step A1 of UnionCom from 4 to 10, the label transfer accuracies of UnionCom are stably around 80% (Fig. 5(b)). When embedding the two dataset into a common space of different dimensionality of *p* in step A3 of UnionCom from 10 to 100, the label transfer accuracies of UnionCom are stable with small fluctuation (Fig. 5)(c). Finally, when choosing different hyperparameter *ρ* for the penalty term Φ(**F**) in Eq. (2), the training loss of the prime-dual algorithm converges very fast to the same value when *ρ* ranges from 5 to 20; however, when *ρ* = 0 which means no penalty term is applied, the loss shows a strong oscillation pattern and convergences slowly (Fig. 5(d)), indicating the necessity of adding the penalty term.

**Figure 5:**
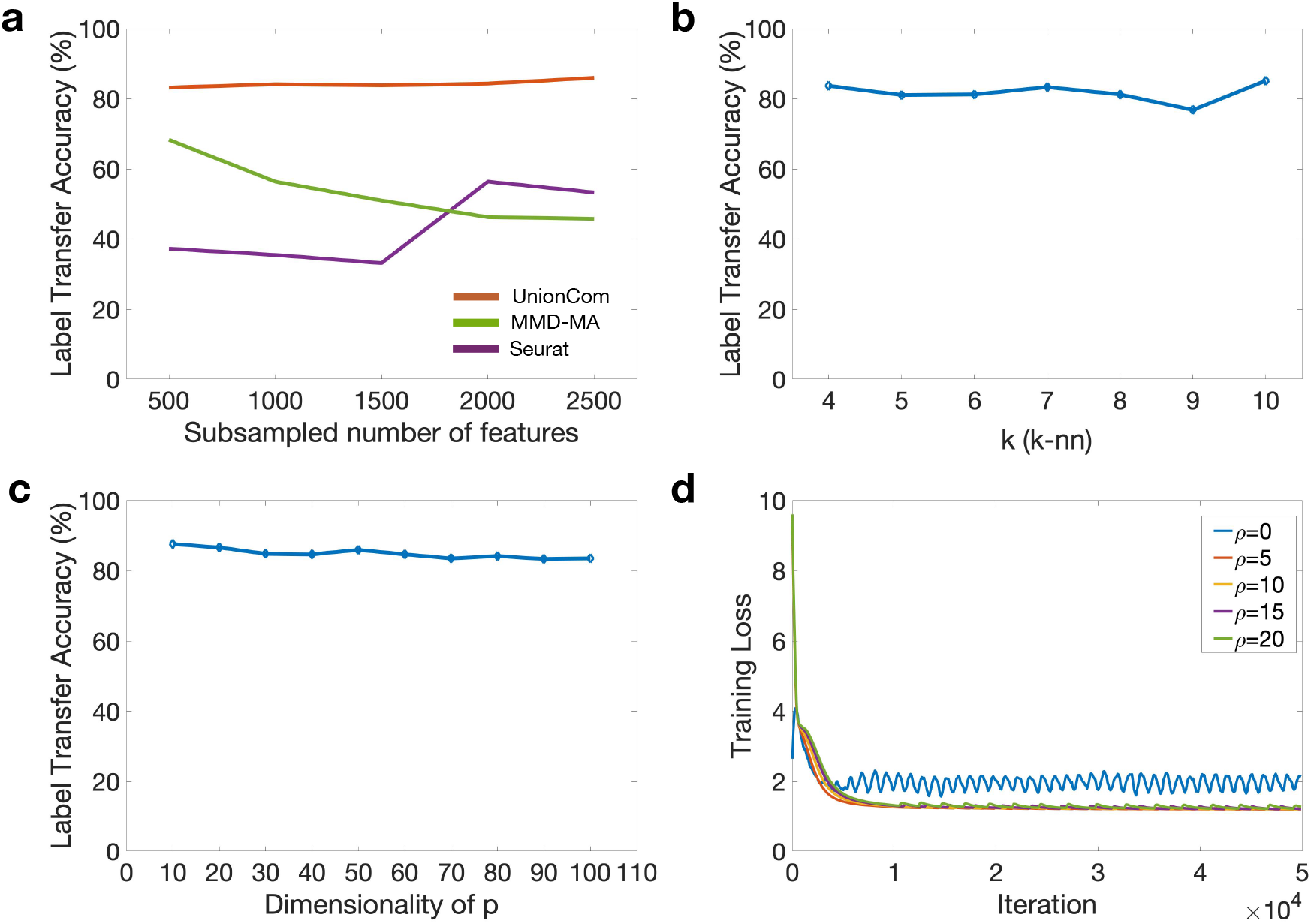
The robustness of UnionCom in both subsampling features of data and parameter choices on the two scNMT datasets of DNA methylation and chromatin accessibility. (a) Label Transfer Accuracy of UnionCom, Seurat, and MMD-MA when randomly sampling a subset of features without replacement from each of the DNA accessibility and chromatin methylation datasets separately before alignment. (b) Label Transfer Accuracy of UnionCom when choosing different *k* of the *k*-nn graph in step A1 of UnionCom. (c) Label Transfer Accuracy of UnionCom when embedding the two datasets into a common space of different dimensionality of *p* in step A3 of UnionCom. (d) The convergence performance of UnionCom in training loss using different *p* for the penalty term in Eq. (2); when *ρ* = 0, no penalty term is applied in Eq. (2).

## 4 Discussion

Manifold alignment is one of the foremost research fields of machine learning and data science [21, 22, 23, 24, 25, 26, 13, 11]. In this study, we develop UnionCom, the unsupervised topological alignment method for single-cell multi-omics data integration. UnionCom represents the intrinsic topological structures embedded in data as distance matrices and then formulates the alignment problem into a convex soft matching problem of matrices. It is totally unsupervised and data-driven, and it does not need the correspondence information, either among cells or among features.

Different from methods such as Seurat and scAlign which conduct the dimensionality reduction prior to the alignment of the cells, UnionCom first aligns the cells across datasets based on the geometrical distance of metric space and then projects the distinct features into a common low-dimensional embedded space. We thus propose a global scaling factor *α* to account for the global geometrical distortions on the embedded intrinsic topological structures and remove the need for pre-matching and normalizing distinct features across multi-omics datasets. Since the un-matched features across datasets have different distributions with complex intercorrelations, normalization of features across single-cell multi-omics datasets to the same distribution can further distort the intrinsic geometric structures, making the alignment problem more difficult. For complex embedded hierarchical structures with multi-scales, UnionCom can align the manifold recursively by introducing scaling-specific factors for each scale of the manifold, and we plan to pursue this topic in our future work.

